# Vagus nerve stimulation limits colonic inflammation through distinct neuroimmune circuitry shaped by inflammatory history

**DOI:** 10.64898/2026.05.28.724702

**Authors:** Kristina Sanchez, Jungjae Park, Emmy Tay, Gatha Pore, Abigail Wagner, Sarah Lee, Jane Li, Ayra Mirza, Melanie Gareau, Colin Reardon

**Author notes:** Corresponding Author: Colin Reardon PhD, Associate Professor, University of California, Davis, 1089 Veterinary Medicine Drive, VM3B, Room 2023, Davis, CA 95616.

## Abstract

Bidirectional communication between the nervous and immune systems has been demonstrated to limit or enhance immune cell function across organ systems and conditions. Although these neuroimmune circuits can become activated as an anti-inflammatory reflex, vagus nerve stimulation **(VNS)** reduces inflammation in models of endotoxemia, rheumatoid arthritis, and intestinal inflammation. In the spleen during endotoxemia, VNS activates a “cholinergic anti-inflammatory pathway” (CAIP), whereby choline acetyltransferase (ChAT)-expressing T cells release acetylcholine to reduce macrophage activation. VNS can also drive CAIP-independent pathways to reduce inflammation in the spleen and intestinal tract, although the circuitry modulating colonic inflammation remains underexplored. Here, we demonstrate that left cervical VNS reduces acute LPS-induced inflammation, evidenced by reduced *Tnfa* expression in colon and spleen and decreased circulating TNFα. In the colon, these protective effects required efferent but not afferent VNS and were independent of ChAT+ T cells, IL10, β-adrenergic signaling, and colonic sympathetic innervation. Critically, the ability of VNS to modulate colonic inflammation depended on prior tissue-specific inflammation. Mice recovering from DSS colitis, despite near-complete histological recovery, were refractory to the protective effects of VNS in the colon. This lack of efficacy in the colon was not reflected in measures of inflammation in the spleen or serum, highlighting the need for target-organ-specific monitoring. This loss of efficacy after colonic inflammation was transient, with restoration occurring upon complete recovery. These findings demonstrate that VNS efficacy in colonic inflammation depends on circuitry distinct from canonical systemic anti-inflammatory pathways, and that tissue responsiveness is shaped by anatomical site and inflammatory history.

**Key Points:**

- Electrical stimulation of the left cervical vagus nerve reduces LPS-induced inflammation in the mouse colon.
- This colonic anti-inflammatory effect requires vagal efferents but not afferent signaling.
- Unlike canonical splenic anti-inflammatory pathways, the colonic response does not require ChAT+ T cells, IL-10, β-adrenergic signaling, or local sympathetic innervation.
- Recent DSS colitis abolishes colonic responsiveness to VNS despite preserved splenic and systemic anti-inflammatory effects.
- Recovery of VNS anti-inflammatory efficacy after colitis shows that neuroimmune responsiveness in the colon is dynamically shaped by inflammatory history.

## Introduction

Sepsis triggers dysregulated inflammatory processes that can extensively damage multiple organs and remains a major cause of mortality (1). As a mucosal tissue that excludes pathogens and other innocuous organisms and substances, this intestinal barrier is a critical determinant of health (2). The gastrointestinal (GI) tract is both a target and amplifier of inflammation (3), as inflammation can reduce intestinal barrier function, allowing translocation of luminal contents, including enteric bacteria. As the GI tract can sustain and worsen sepsis-associated pathology, modulation of intestinal inflammation may have therapeutic value and limit the development of further sequelae. Although many cell types respond to bacterially derived pathogen-associated molecular patterns such as Lipopolysaccharide (LPS), specific subsets of macrophages adjacent to intestinal epithelial cells have been identified as critical responders, producing proinflammatory cytokines such as TNFα (4). Despite this knowledge, approaches that effectively and selectively suppress the causes of inflammation and, consequently, damage during sepsis remain limited. Immune cell action can be positively and negatively regulated through bidirectional communication with the nervous system (5). Specialized neurons detect infection and inflammation and can shape the immune cell function in diverse organs, including the lung, lymph nodes, spleen, and GI tract. One of the best characterized neuroimmune pathways that inhibits inflammation during endotoxemia is the cholinergic anti-inflammatory pathway (CAIP), in which activation of vagal afferent neurons (sensory) due to inflammation activates vagal efferent signaling to drive sympathetic innervation of the spleen and MLN to activate a highly specialized population of T-cells (6–10). These CD4+ T-cells express the enzyme choline acetyltransferase (ChAT), enabling them with the capacity to produce and release Acetylcholine (ACh) in response to β2-adrenergic receptor (β2AR) activation (9). These cells are critical to CAIP, as released ACh suppresses pro-inflammatory cytokine production by acting on nicotinic α7 receptors (α7R nAChR) (6–9, 11) and inhibiting NF-κB signaling (5). Additional neuroimmune circuits with overlapping functionality have also been discovered to inhibit LPS-induced inflammatory responses in the spleen, and in the lung, by vagal afferent activation of the sympathetic nervous system (12, 13). Highlighting the multitude of neuroimmune circuits in distinct segments of the intestinal tract, vagal efferent stimulation reduced post-operative ileitis by reducing macrophage activation independently of α7nAChR (14). Whether VNS can regulate immune cells within the colon remains unclear, particularly given that the colon has relatively sparse, though not absent, vagal innervation and extensive intrinsic innervation within the enteric nervous system (ENS) (15, 16).

A major limitation of prior studies using neural stimulation to abrogate inflammation is the nearly exclusive use of previously healthy subjects. Neuroimmune circuits are comprised of biological components in tissues shaped by prior acute or chronic inflammation, yet the consequence of this inflammatory history, and how this alters responsiveness to neuronal stimulation, is poorly understood. This question is especially relevant where clinical trials of VNS in inflammatory bowel disease (IBD) have often failed to demonstrate efficacy, or reach primary clinical endpoints (17).

Here we assess the neuroimmune circuitry elicited by VNS in the context of an acute inflammatory stimulus. Electrical stimulation of the left cervical vagus nerve attenuated LPS-induced colonic inflammation. This protection is afforded by efferent but not afferent VNS. Our studies further identified that β-adrenergic receptors were not required and that VNS-mediated suppression in the colon is independent of sympathetic innervation. Highlighting that our observations are distinct from prior studies, IL10 was not required, and CAIP was excluded as VNS-induced control of inflammation was unaffected in ChAT T-cell conditional knockout mice.

The influence of prior inflammation on responsiveness to VNS was determined using the well-established dextran sodium sulfate (DSS) model of transient self-resolving colitis (18). In mice that recovered from colitis at day 14 post-DSS, VNS no longer reduced LPS-induced inflammation despite preserved anti-inflammatory effects in the spleen, and reduced TNFα detected in serum. After a longer recovery period, colonic responsiveness to VNS was restored, suggesting that prior inflammation transiently alters the capacity of target tissues to respond to neural anti-inflammatory signals.

Together, these data indicate that selective VNS can control endotoxemia-induced colonic inflammation through a discrete neuroimmune circuitry and highlight that this protection may be temporarily lost in tissues with recent inflammation. More broadly, they suggest that a history of inflammation may influence responsiveness to bioelectronic therapies and should be considered when designing or applying neurostimulation-based interventions.

## Results

### Vagus nerve stimulation suppresses colonic inflammation during endotoxemia

To evaluate the effect of VNS on the inflammation that occurs during endotoxemia in the intestinal tract, mice were subjected to sham or electrical VNS to the left cervical vagus for 10 minutes before injection of PBS (vehicle) or LPS, with VNS continued for a total of 20 minutes **(Figure 1A)**. Colonic inflammation induced by LPS injection (4 mg/kg i.v) was reduced by VNS but not in non-stimulated controls, as indicated by a significant reduction in the expression of *Tnfa*, *Ccl4*, and *Ccl2* mRNA **(Figure 1B)**. VNS did not prevent the expression of all LPS-induced genes in the colon, as *Cxcl1* was not significantly reduced compared to non-stimulated control subjects. Although LPS increased IL10 in the colon, this expression was not further enhanced in VNS-treated subjects compared to sham controls (**Figure S1A**). The anti-inflammatory effects of left cervical VNS were not restricted to the colon, as *Tnfa* expression in the spleen was also reduced **(Figure 1C)**. These reductions in proinflammatory cytokines occurred in the absence of increased *Il10* expression, as well as expression of enzymes that produce anti-inflammatory lipid mediators (**Figure S1A).**

**Figure 1.**
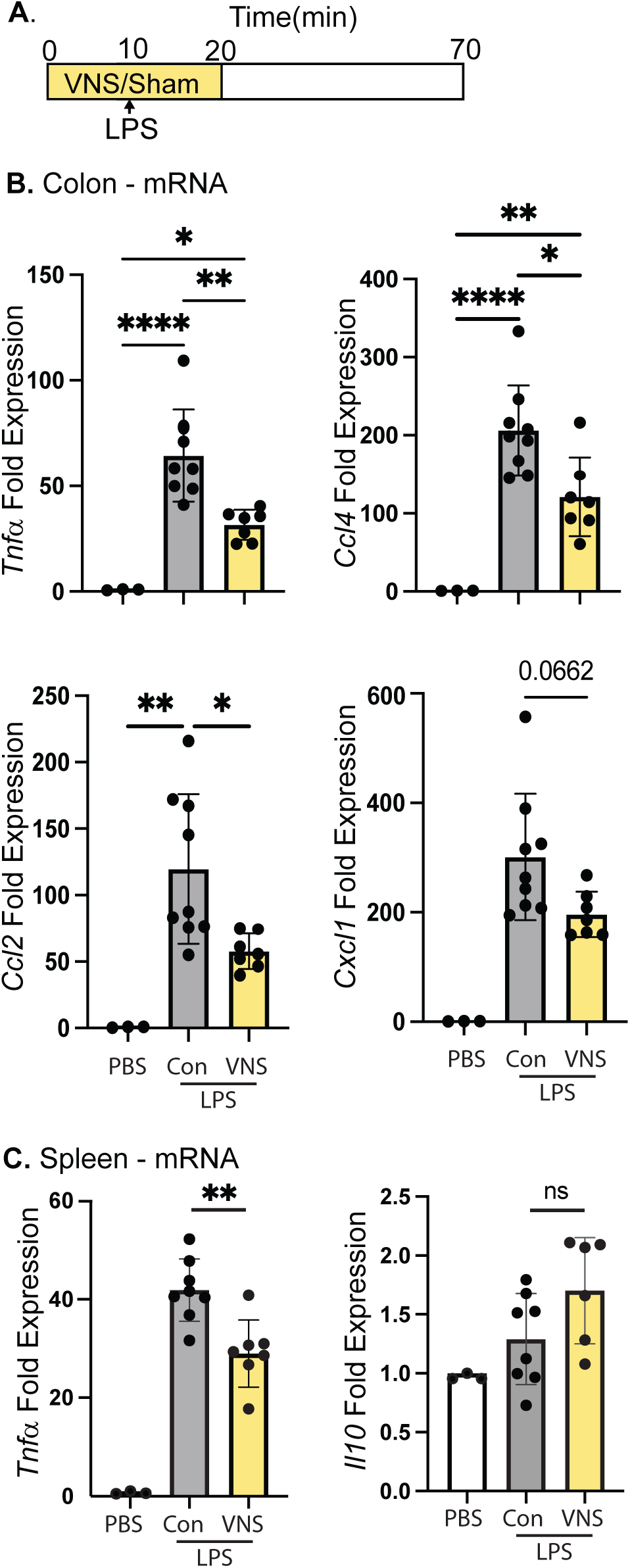
Vagus nerve stimulation suppresses colonic inflammation during endotoxemia. **(A)** Schematic of the acute endotoxemia model in which LPS (4 mg/kg i.v) is administered after the first 10 minutes of VNS. Colon and spleen were collected for qRT-PCR analysis. **(B)** Colonic mRNA expression of *Tnfα, Ccl4, Ccl2, and Cxcl1,* and **(C)** splenic mRNA expression of *Tnfα* and *Il10*. Data are presented as mean ± SD. Statistical analysis was performed using one-way ANOVA with Tukey’s multiple comparisons test. *p< 0.05; **p<0.01; ***p<0.001; ****p<0.0001.

### VNS control of colonic inflammation during endotoxemia is independent of IL10 and CAIP

To definitively assess the role of IL10 in the anti-inflammatory circuit, we performed electrical VNS in WT and *IL10^−/−^* mice. VNS reduced LPS-induced expression of colonic *Tnfa* and *Ccl4* in both WT and *IL10^−/−^* mice compared to controls that received LPS alone **(Figure 2A & B)**. Although VNS reduced *Icam1* expression in WT, no attenuation of LPS-induced expression was observed in *IL10^−/−^*mice **(Figure 2C)**, and *Vcam1* expression was only reduced in VNS-treated *IL10^−/−^* mice **(Figure 2D)**.

**Figure 2.**
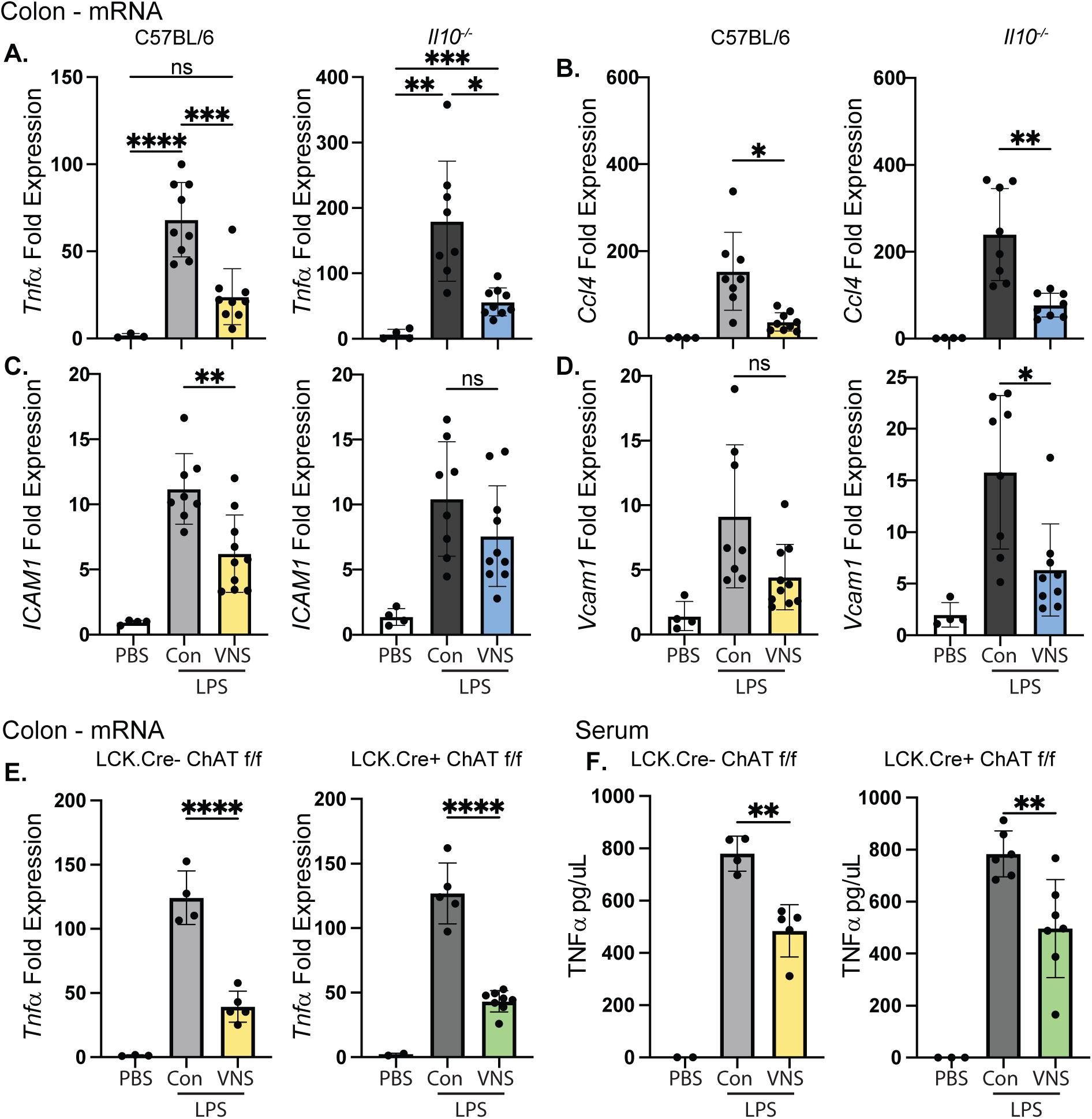
VNS-mediated control of colonic inflammation during endotoxemia is independent of IL10 and CAIP. Colonic mRNA expression of **(A)** *Tnfα*, **(B)** *Ccl4*, **(C)** *Icam1*, **(D)** *Vcam1* in WT vs *Il10^−/−^*mice after control, and LPS ± VNS. **(E)** Colonic mRNA *Tnfα* and **(F)** serum TNFα in WT (LCK.Cre-ChATf/f) and ChAT T cell cKO (LCK.Cre+ ChATf/f) mice. Data are presented as mean ± SD. One-way ANOVA and post-hoc analysis with Tukey’s multiple comparisons test were used for statistical analysis. *p< 0.05; **p<0.01; ***p<0.001; ****p<0.0001.

To assess the putative contribution of the cholinergic anti-inflammatory pathway, we used wild-type or T-cell-conditional knockouts of Choline acetyltransferase (ChAT). VNS was applied to the left cervical vagus nerve in WT (LCK.Cre^−^ ChAT^f/f^) and ChAT T-cell cKO (LCK.Cre^+^ ChAT^f/f^) mice. Colonic inflammation induced by LPS in WT mice and ChAT T-cell cKO mice was significantly reduced with VNS (**Figure 2E)**. Systemic inflammation, indicated by serum TNFα protein, was also significantly reduced by left cervical VNS **(Figure 2F)**, suggesting that left VNS reduces systemic and colonic inflammation independently of CAIP.

### VNS-mediated control of colonic inflammation requires vagal efferents but not colonic sympathetic neurons

To evaluate the effect of afferent and efferent VNS on intestinal inflammation during endotoxemia, the left cervical vagus nerve was transected immediately prior to VNS **(Figure 3A)**. Efferent VNS, but not afferent VNS, elicited a reduction of colonic inflammation **(Figure 3B).** Antagonism of pan-β-adrenergic receptors prior to VNS revealed that these receptors are not required for the suppression of *Tnfa* in the colon **(Figure 3C)**. To assess the contribution of colonic sympathetic neurons during VNS, we performed a colon-specific sympathectomy via intrarectal administration of 6-OHDA 3 days prior to LPS challenge and VNS **(Figure 3D).** Colonic inflammation, indicated by *<u>Tnfa</u>* and *Cxcl1* mRNA expression induced by LPS in vehicle and 6-OHDA-treated mice, was reduced with VNS **(Figure 3E & F)**. Colonic selective sympathectomy was confirmed via reduction of *Th* mRNA in colon but not spleen **(Figure 3G & J).** Systemic *Tnfa* but not *Cxcl1* mRNA expression was controlled with VNS in either sham or 6-OHDA-treated mice **(Figure 3H & I).**

**Figure 3.**
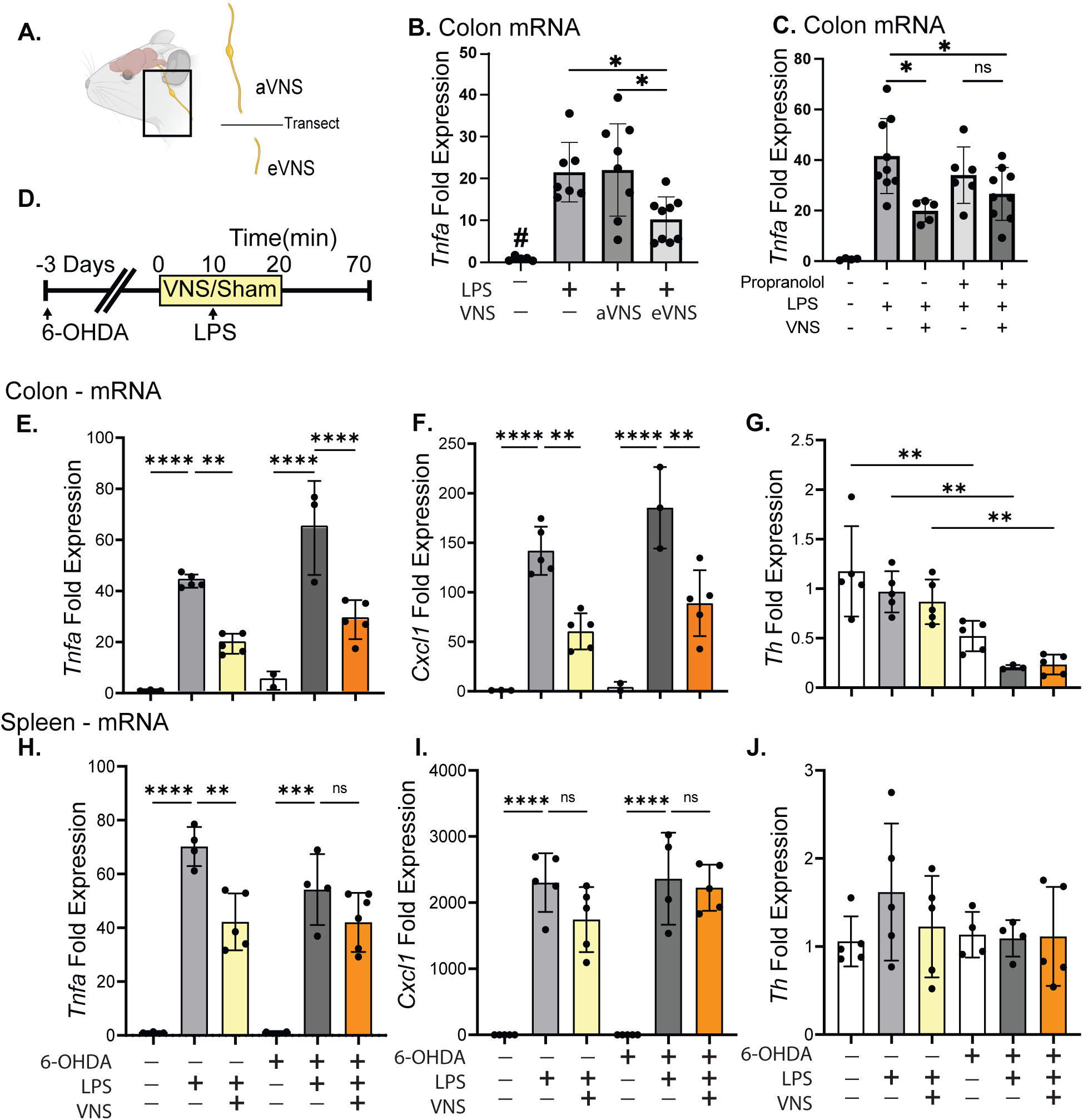
VNS suppresses colonic inflammation through vagal efferents independently of colonic sympathetic neurons. **(A)** Experimental schematic for selective afferent or efferent VNS. **(B)** Colonic *Tnfα* mRNA expression after afferent VNS (aVNS) or efferent VNS (eVNS). (C) Colonic *Tnfα* mRNA expression in mice receiving vehicle or propranolol prior to LPS challenge and VNS **(D)** Experimental schematic for colonic sympathectomy by intrarectal 6-OHDA administration 3 days prior to LPS challenge with or without intact VNS. **(E)** Colonic mRNA of *Tnfα* **(F)** *Cxcl1* and **(G)** *Th.* **(H)** Splenic mRNA expression of *Tnfα* **(I)** mRNA of *Cxcl1* and **(J)** Th. Data are presented as mean ± SD. One-way ANOVA and post-hoc analysis with Tukey’s multiple comparisons test were used for statistical analysis. *p< 0.05; **p<0.01; ***p<0.001; ****p<0.0001.

### Prior colonic inflammation transiently disrupts responsiveness to VNS

To determine whether prior colitis reduces the efficacy of VNS, mice were subjected to DSS-induced colitis and randomized to receive PBS, LPS (sham), or LPS with VNS **(Figure 4A)**. DSS induced a mild, transient colitis that peaked at Day 8, and largely resolved by Day 14, as determined by body weight loss and histopathology **(Figure 4B & C)**. Although overt inflammation was no longer evident, 14 days post-DSS, VNS was ineffective in the colon compared to naïve controls **(Figure 4B)**. Despite this loss of efficacy in the colon, VNS remained effective in the spleen **(Figure 4C)** and reduced serum TNFα **(Figure 4D)**. Given that colonic inflammation can negatively affect the ENS, we sought to determine whether DSS treatment and recovery altered the number of colonic myenteric neurons. Neuron counts were comparable between sham and DSS-treated mice, regardless of recovery window (**Figure 5A & B**). Thus, the loss of VNS efficacy after DSS was not explained simply by the loss of myenteric neurons.

**Figure 4.**
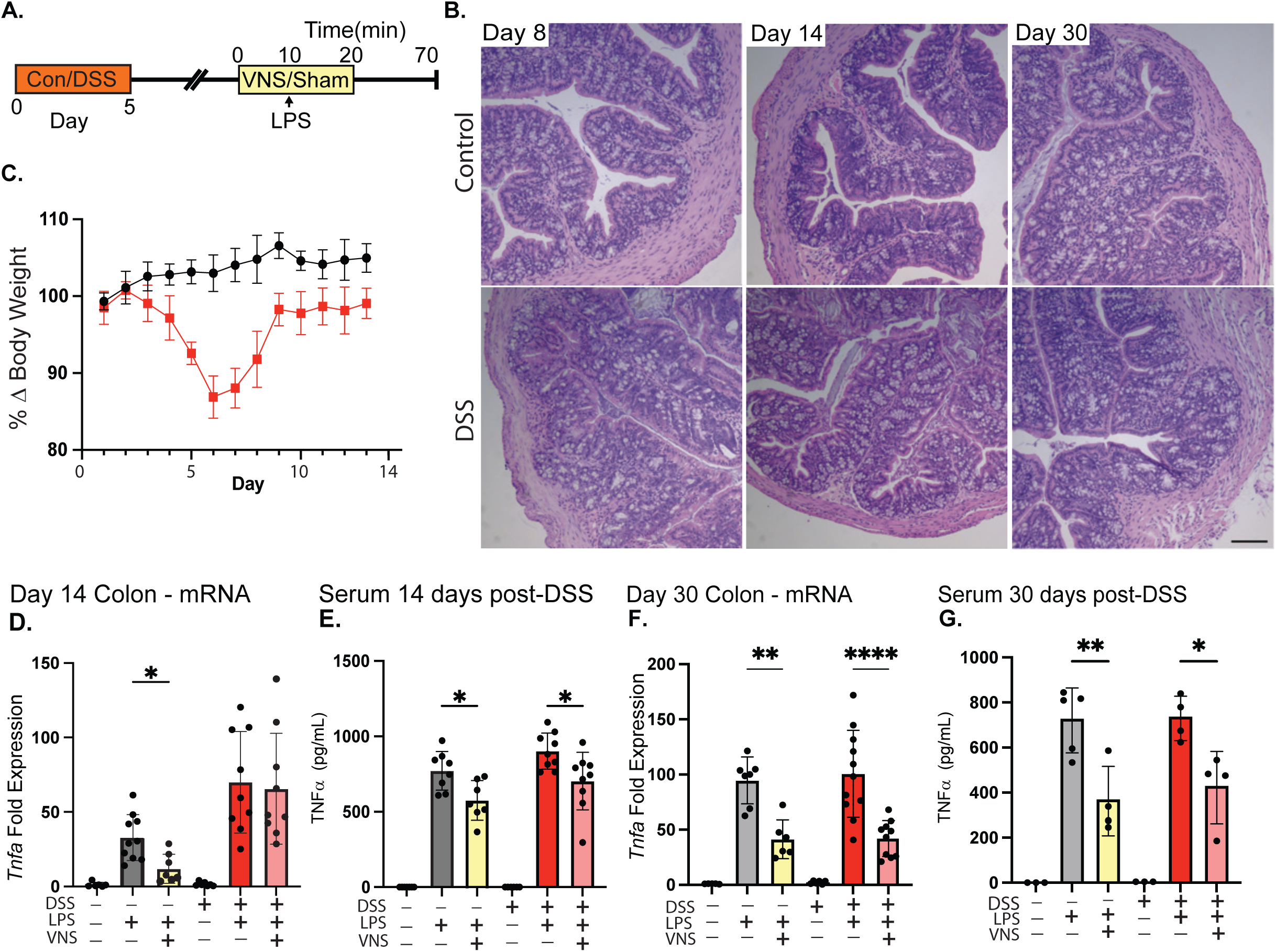
Prior colonic inflammation transiently disrupts responsiveness to VNS. **(A)** Experimental schematic of DSS colitis followed by acute LPS challenge 14 days post-DSS. **(B)** Representative mid-distal colon histology on days 8, 14 and 30 post-DSS administration. Scale bar indicates 100 µm. **(C)** Body weight throughout DSS treatment, peak inflammation, and recovery. **(D)** Colonic *Tnfα* mRNA expression, and **(E)** serum for TNFα at day 14 post-DSS. **(F)** Colonic *Tnfα* mRNA expression, and **(G)** serum for TNFα at day 30 post-DSS. Data are presented as mean ± SD. One-way ANOVA with Tukey’s multiple comparisons test. *p< 0.05; **p<0.01; ***p<0.001; ****p<0.0001.

**Figure 5.**
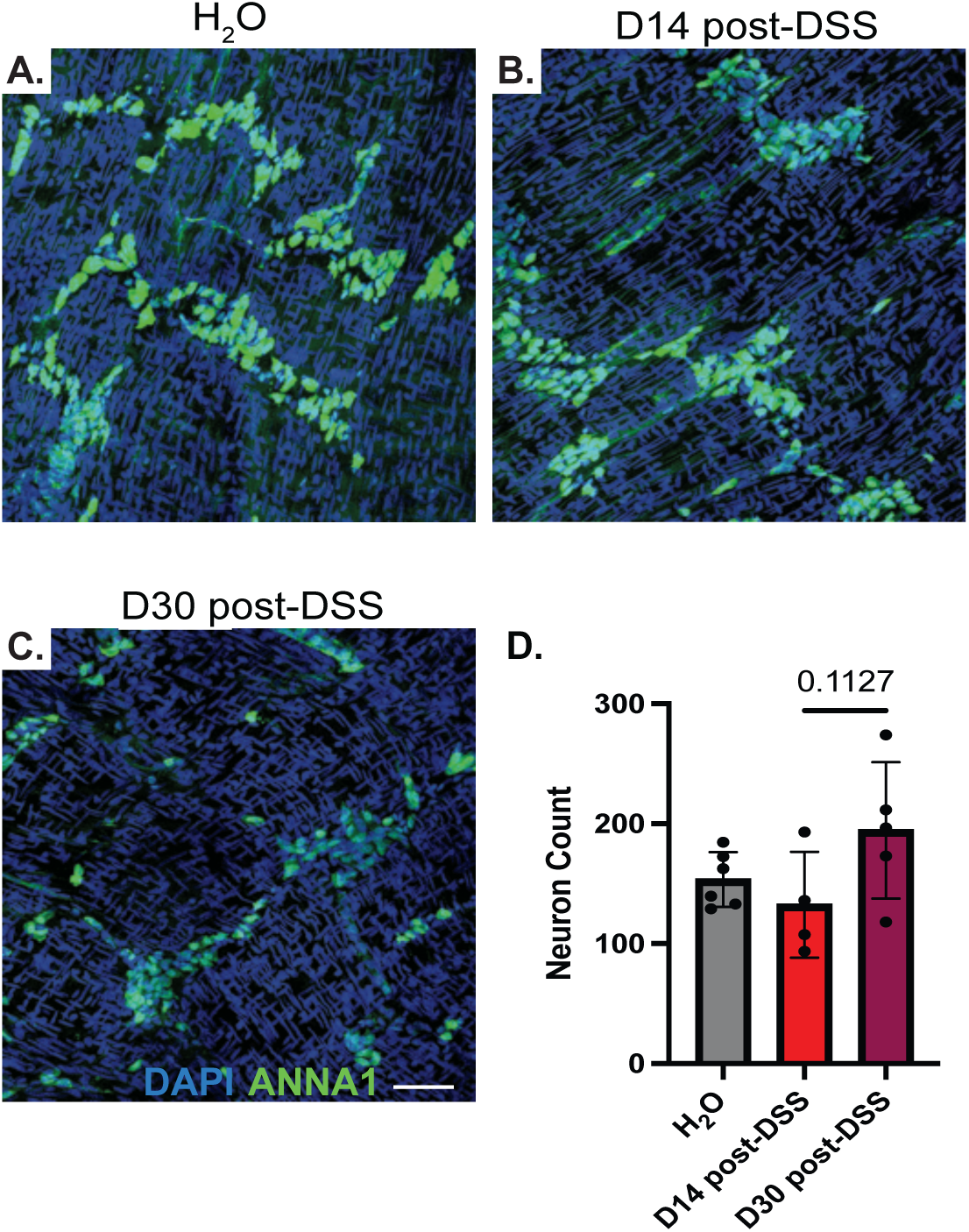
Prior colonic inflammation does not impact total number of enteric neurons. **(A)** Whole-mount staining of enteric neurons (ANNA1) in colon from sham, Day 14-post-DSS, and Day 30-post-DSS-treated mice. Scale bar indicates 50 µm. **(B)** Neuron counts of enteric neurons post-DSS. Data are presented as mean ± SD. One-way ANOVA and post-hoc analysis with Tukey’s multiple comparisons test were used for statistical analysis.

### Prior infection of the ileum does not affect the ability of VNS to reduce inflammation

To determine whether the loss of VNS efficacy observed after DSS colitis could be generalized to other forms of intestinal inflammation, we used neonatal enteropathogenic *E. coli* infection and challenged adult mice with LPS with or without VNS. Neonatal mice were infected at P7 (**Figure 6A**), and bacterial burden was monitored weekly until clearance at P28 (**Figure 6B**). As EPEC primarily infects the ileum, the ability of VNS to down-regulate inflammation in this region was assessed following LPS administration. In addition to the expected reduction in serum TNFα (**Figure 6C**), VNS significantly reduced LPS-induced ileal inflammation in sham-infected and neonatally EPEC-infected mice, as indicated by a significant reduction in the expression of *Tnfa, Ccl2, Ccl4, and Cxcl1* mRNA (**Figure 6D**). These findings indicate that prior intestinal inflammation does not uniformly disrupt VNS-induced anti-inflammatory effects and that the impact of prior inflammation depends on the tissue, timing, and nature of prior insult.

**Figure 6.**
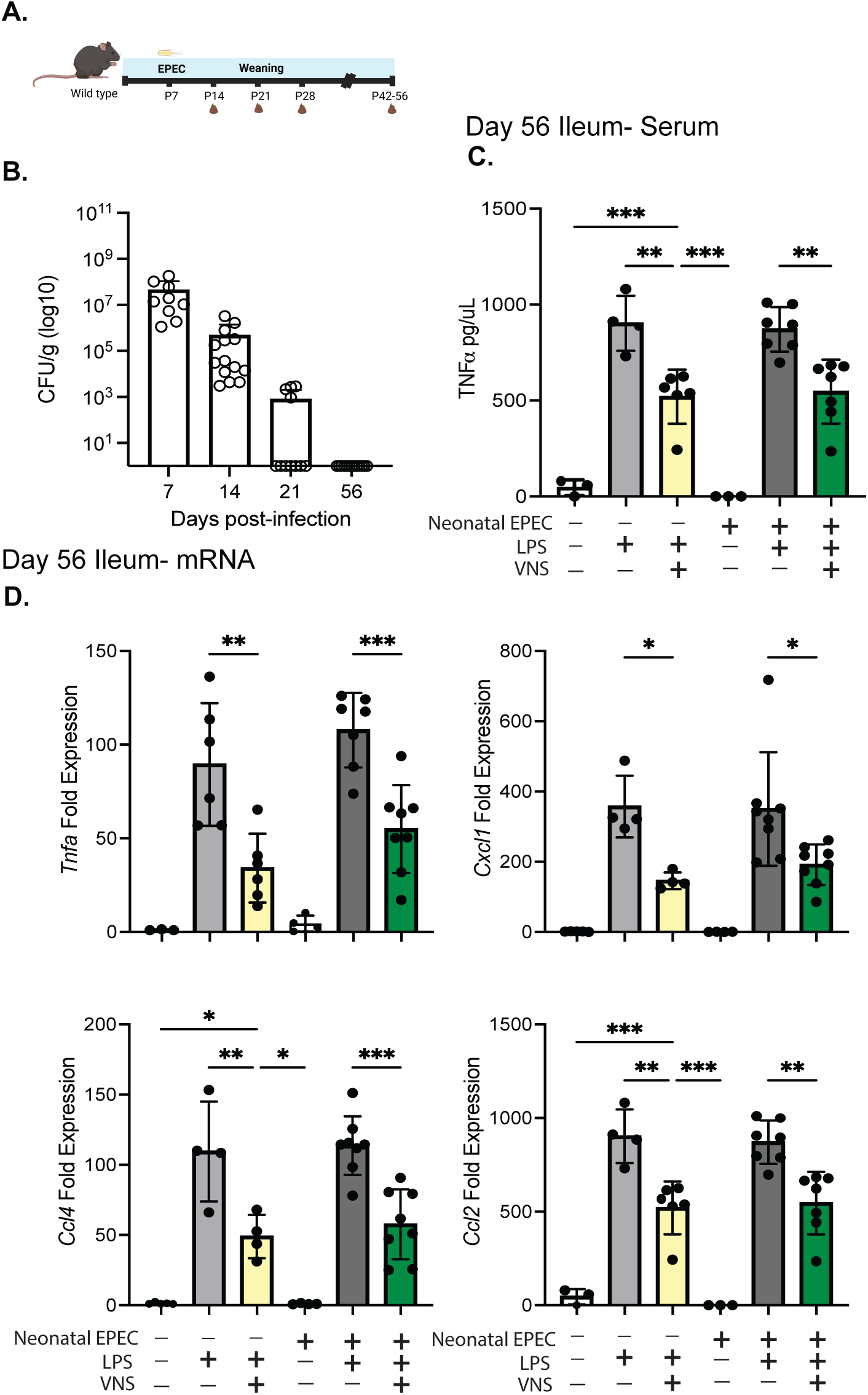
Prior infection of ileum does not affect the ability of VNS to reduce inflammation. **(A)** Schema of murine neonatal infection model prior to acute challenge of LPS on Day 56 post-infection. Mice were gavaged with enteropathogenic *E. coli* at p7 and infection level was monitored weekly until clearance. **(B)** CFU of fecal samples throughout infection timeline. **(C)** Serum for TNFα Day 56 post-EPEC infection, **(D)** mRNA for *Tnfα* in Day 56 post-EPEC infection in ileum. Data are presented as mean ± SD. One-way ANOVA and post-hoc analysis with Tukey’s multiple comparisons test were used for statistical analysis. *p< 0.05; **p<0.01; ***p<0.001; ****p<0.0001.

## Discussion

Inappropriate or excessive immune cell activation and its sequelae are critical aspects of many pathologies, including septic shock (1). Although many organ systems are affected during sepsis-induced inflammation, the gastrointestinal tract acts as both a target and a contributor to disease severity (3). Elegant studies have demonstrated that several neuroimmune circuits can restrain this inflammation and reduce immunopathology (9, 19–22). These neuroimmune circuits can be activated either as part of a reflex arc or by providing exogenous neuronal stimulation. Optogenetic stimulation targeting ChAT+ enteric neurons has been shown to improve DSS-associated immunopathology, and ablation of this subset of enteric neurons resulted in more severe colitis (23). Complementary work has also demonstrated that enteric neuronal Piexo1-dependent mechanosensing maintains both mechanical and immunological homeostasis, providing additional evidence that defined enteric neuronal programs modulate colitis severity (24). Electrical stimulation of the right vagus nerve has been shown to reduce small intestinal macrophage activation during post-operative ileitis, whereby nicotinic α4β2 receptors were required for VNS-induced control of inflammation (14). Other approaches, including abdominal VNS, were also shown to be effective in reducing proinflammatory serum cytokines and in reducing the recruitment of immune cells to the small intestine (25). Although these data support the ability of VNS to reduce intestinal inflammation, vagal innervation is not consistent throughout the GI tract. Vagal innervation of the colon is present but significantly reduced compared with that of the small intestine (26); the ability of this innervation to influence immune outcomes was unknown. Despite these studies, the ability to control intestinal inflammation during endotoxemia has not been previously demonstrated.

We demonstrate that electrical VNS to the left cervical vagus reduced inflammation in the colon and spleen during endotoxemia. Given the previously identified induction of IL10 by VNS as the suggested mechanism of inflammatory control (27), we assessed the role of this cytokine by using a global genetic knockout. VNS did not increase *IL10* mRNA expression in the colon or spleen compared to controls, and VNS efficacy was not reduced in *IL10^−/−^* mice. These data indicate that the reduction of endotoxemia-induced inflammation achieved by left cervical VNS occurs independently of the anti-inflammatory cytokine IL10. Since it is well established that stimulation of the right cervical vagus nerve induces the CAIP, we assessed the potential role of this pathway in regulating endotoxemia-induced colonic inflammation after stimulation of the left vagus nerve (6, 9). Mice with T-cell conditional knockout of ChAT, and therefore deficient in the key mechanism of CAIP (9), and WT controls were equally protected by left VNS. This is consistent with the exceptional rarity of these ChAT+ T-cells in the colon of naïve mice, as we have shown previously (18). We and others have also identified CAIP-independent pathways whereby right cervical afferent VNS activated sympathetic innervation of the spleen and MLN to reduce inflammation (19, 28). Here, we have identified that efferent but not afferent left VNS is sufficient to reduce colonic inflammation. These data, in combination with the independence from colonic sympathetic innervation and β-adrenergic receptors, demonstrate that a vagal afferent to sympathetic nervous system pathway is not required. Our data argue against a simple extension of known anti-inflammatory reflexes in the colon and favor engagement of a different peripheral effector pathway.

This regulation of inflammation was disrupted after recent DSS-induced colonic inflammation but not neonatal enteric bacterial infection of the small intestine, suggesting that the impact of inflammatory history on VNS responsiveness depends on the timing, location, and nature of the prior insult. Consistent with this idea, DSS is known to remodel the ENS, including enteric glia, while also recruiting activated immune cells and altering epithelial and stromal tissue architecture (29–32). Prior work further demonstrates altered expression of neurotransmitter receptors and modulators of intracellular signaling, including phosphodiesterase (PDE)-regulated cAMP signaling, which may influence cellular responses to neuronal inputs (33, 34). Thus, DSS colitis is associated not only with changes in the cellular composition and tissue architecture of the colon but also with altered responsiveness to select inputs. Recent work showing altered vagal afferent sensory encoding following DSS colitis supports the broader concept that intestinal inflammation can reshape neuroimmune communication. DSS-induced inflammation significantly increased the number of spontaneously active vagal afferents, but with significantly attenuated signal amplitude and evoked responses to stimuli (35). This supports the contention that colonic inflammation can impinge on anti-inflammatory neuroimmune circuits. However, our findings remain distinct because the colonic response required efferent VNS, and prior DSS colitis selectively disrupted efficacy in the colon while leaving serum responses intact. Thus, prior inflammation did not eliminate the ability of VNS to generate an anti-inflammatory response but instead impaired the ability of the previously inflamed target tissue to respond to that stimulation. This distinction may be important for understanding both variable responses to VNS and why systemic anti-inflammatory effects are not always mirrored in the affected organ.

Despite many preclinical successes in sepsis and other inflammatory conditions, as well as recent clinical studies in rheumatoid arthritis, the efficacy of VNS for intestinal inflammation remains unclear (16, 20, 36, 37). Reduced inflammation and improvements in colitis disease activity index have been observed in a small open-label clinical study, but responses are variable, and patients experiencing disease flares prior to or during study enrollment appear to derive less benefit (16, 17). Similar anecdotal observations have been made in non-human primates treated with VNS for spontaneously developing intestinal inflammation (38). Our findings provide experimental support for this idea by demonstrating that recent colonic inflammation can transiently impair the local anti-inflammatory effect of VNS, even when systemic anti-inflammatory effects are preserved.

The mechanism underlying this transient loss of local colonic responsiveness remains unresolved. DSS colitis can alter tissue architecture, replace resident macrophages with newly recruited monocytes, and reprogram tissue resident populations that may be required to relay VNS-induced signals. Repeated bouts of intestinal inflammation induces CCL2-dependent monocyte recruitment into the myenteric plexus, contributing to neuronal loss, ENS remodeling, and consequently dysmotility (30). Although the number of colonic myenteric plexus neurons was not reduced, this does not exclude inflammation-induced changes in neuronal composition, neurotransmitter production, or receptor expression, glial activity, or neuroimmune interactions (39, 40). Thus, the post-DSS colon may retain a normal neuronal architecture while temporarily losing the functional capacity to respond to VNS, consistent with the restoration of effect with a prolonged recovery period. The neonatal EPEC model provided critical insight into the role of inflammatory history in VNS responsiveness. Neonatal infection was used to determine if early-life intestinal inflammation produces long-term neuroimmune alterations that impair the protective effects of VNS later in life. Although neonatal EPEC infection can induce ileal inflammation, VNS still reduced endotoxemia-induced ileal inflammation in adulthood. These findings suggest that prior inflammation does not uniformly impair VNS efficacy, but that its impact depends on the nature of the insult, timing, location, and recovery state. More broadly, these findings suggest that inflammatory state may strongly influence clinical efficacy and that systemic biomarkers may be insufficient to assess therapeutic success.

## Methods

### Animals

C57BL/6J, IL10^−/−,^ and LCK.Cre+ were purchased from Jackson Laboratories and used to establish a breeding colony at UC Davis. Mice possessing the ChAT allele flanked by loxP sites, ChATf/f, have been previously described and crossed with LCK.Cre+ line to produce LCK.Cre+ ChATf/f progeny (13, 18). Male and female mice aged 7-12 weeks were used for experiments. All animals had *ad libitum* access to food and water, and all procedures were approved by the UC Davis Institutional Animal Care and Use Committee (IACUC).

### DSS Administration

Once mice reached a minimum body weight of 18g, 5-day administration of 3% dextran sodium sulfate (DSS) in drinking water began, as previously described (41, 42). Water consumption was measured daily during DSS administration. On day 6, mice were switched back to normal drinking water until the collection timepoint. Body weights were monitored daily throughout the experiment, and mice that lost more than 20% of their initial body weight were euthanized per IACUC guidelines.

### Colonic sympathectomy

Colonic sympathectomy was induced by a single dose of 6-hydroxydopamine (6-OHDA). 6-OHDA was dissolved in 0.2% ascorbic acid and 50% ethanol, then administered at a dose of 100 mg/kg. Briefly, mice were anesthetized, and a flexible tube was inserted intrarectally to a depth of 3 cm from the anus, allowing 100 μL of the 6OHDA solution or 0.2% ascorbic acid/50% ethanol solution (vehicle control) to be slowly administered into the colon. Mice were returned to their cages to recover for three days. Vehicle- and 6OHDA-treated mice were randomly divided into experimental groups for VNS. Colon and spleen tissues were collected to assess sympathectomy and inflammation.

### Vagus nerve stimulation

Mice were anesthetized using 1.5-2% isoflurane, and body temperature was maintained at 37°C using a thermocouple-controlled heating pad. The left cervical vagus nerve was exposed. In select studies, the exposed vagus nerve was left intact, or a small suture was tied before cutting for afferent/efferent stimulation and placed on a bipolar hook electrode (FHC, Bowdoin, ME, USA). Electrical stimulation was conducted via a Grass stimulator S88 with a stimulator isolation unit for 20 minutes (5 V, 2ms, monophasic square wave, 5 Hz), and lipopolysaccharide was injected (LPS, InvivoGen, tlrl-3pelps, 4mg/kg i.v.) after the first 10 minutes of stimulation. Mice were euthanized one-hour post-LPS injection via cardiac puncture and subsequent cervical dislocation. Serum, colon, spleen, and ileum were collected for qRT-PCR analysis.

### Quantitative real-time PCR

Colon and spleen samples were collected for quantitative real time PCR (qRT-PCR) in TRIzol (Invitrogen) and stored at −80°C before further analysis. RNA was extracted by homogenizing tissue using 5 mm stainless steel beads in a TissueLyser (Qiagen). The iSCRIPT reverse transcriptase kit (Bio-Rad) was used to generate cDNA for qPCR analysis on a QuantStudio 6 (Thermo Fisher Scientific, Waltham, MA, USA). The following primer pairs were sourced from Primerbank (43) for amplification:

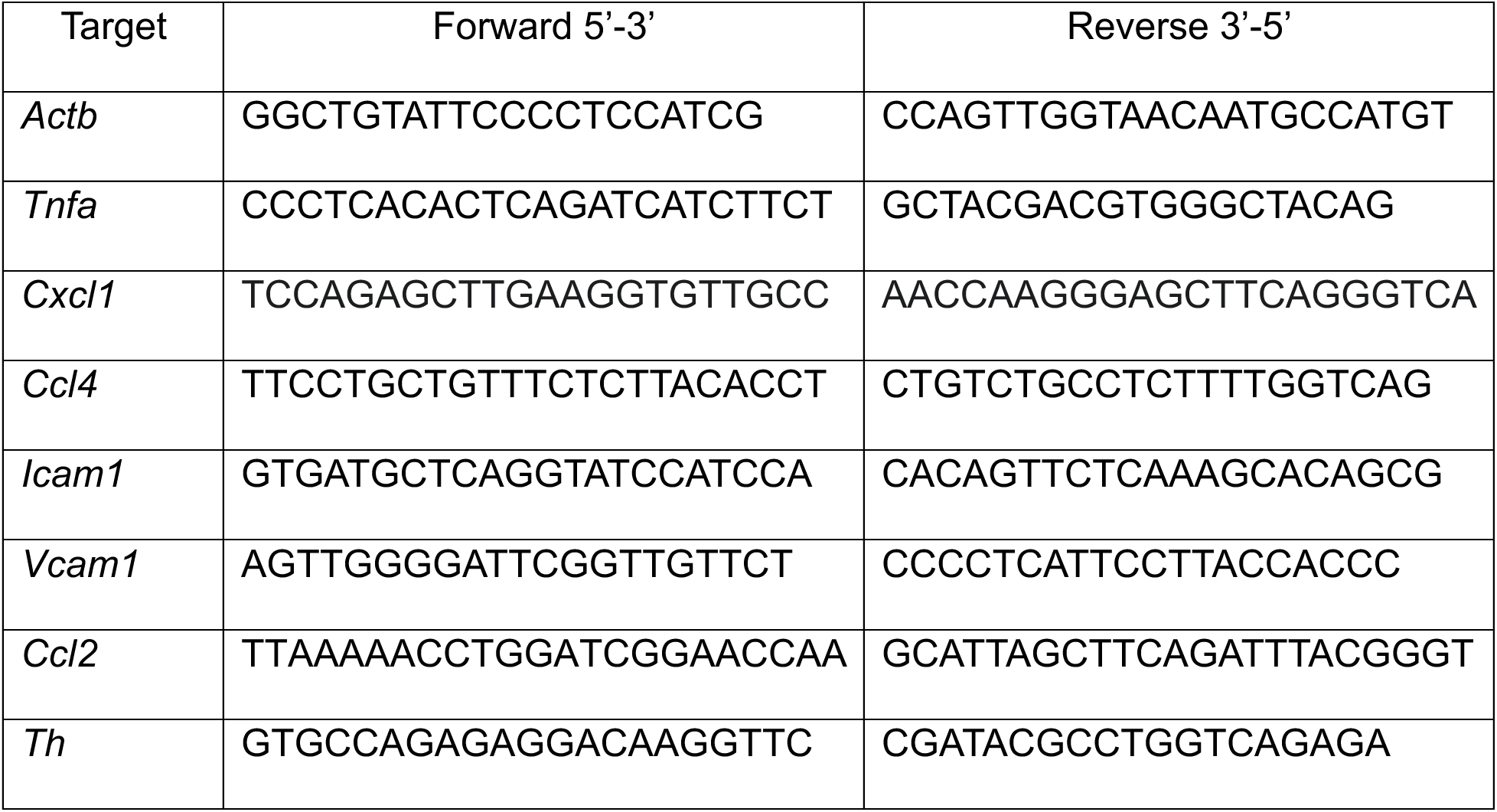

### ELISA

Serum collected by cardiac puncture was isolated using serum separator tubes (BD Biosciences) and frozen at −80°C until assay. Briefly, a sandwich ELISA for TNFα (Thermo Fisher Scientific) was performed by coating a 96-well plate with capture antibody overnight. Serum samples were added and incubated for two hours, and washed thoroughly. The biotin-conjugated detection antibody was added, followed by washing, and the addition of streptavidin HRP. After incubation and final washing, TMB substrate was added. The reaction was quenched after 15 minutes by the addition of 1 N H_2_SO, absorbance at 450 nm and 570 nm was determined using a Synergy H1 plate reader (BioTek Instruments Inc.), and sample concentrations were calculated by linear regression from a standard curve.

### Histology

Tissues were fixed in 10% normal buffered formalin before processing and paraffin embedding. Slides with 6 µm sections were deparaffinized, rehydrated, and stained with Harris hematoxylin according to standard protocols (44). Stained sections were imaged using a Nikon Labophot microscope (Nikon) equipped with a Teledyne BlackFly color camera and using Spinview software (Teledyne Vision Solutions).

### Whole-mount immunostaining of colonic myenteric neurons

Distal colon samples from adult 10–12-week-old mice that had been on DSS for 14 or 30 days were placed in cold, oxygenated PBS (1X), opened along the mesenteric border, and gently rinsed in cold PBS to remove luminal contents. We followed a previously published method for isolation and staining protocols(45). For preparation of colonic whole mounts containing the myenteric plexus, tissues were pinned mucosal side up in a Sylgard-coated dish under a dissecting microscope. The mucosal layer, including the epithelium and lamina propria, was carefully removed using fine forceps and micro-scissors, leaving the submucosa and the longitudinal and circular smooth muscle layers. Although the submucosal layer was retained during tissue preparation, imaging and quantification in the present study were restricted to ANNA-1-positive (human anti-HuC/HuD) myenteric neurons, and submucosal neurons were not included in the analysis. Isolated tissues were fixed in 4% paraformaldehyde (PFA) for 2 h at 4°C, followed by 3 × 15-min washes in PBS. Tissues were blocked and permeabilized overnight at 4°C on a shaker in PBS containing 5% normal goat serum (NGS), 10% bovine serum albumin (BSA), and 0.3% Triton X-100. Neurons were stained with ANNA-1, diluted in blocking buffer, and incubated for 96 h at 4°C with gentle rocking. Tissues were subsequently washed in PBS containing 0.1% Triton X-100 (1 × 5-min wash, followed by 3 × 30-min washes) at 4°C. Secondary antibody was added (goat anti-human IgG [H+L] conjugated to Alexa Fluor 647 diluted in blocking buffer; A21445 [Invitrogen]) and incubated for 24 h at 4°C, and nuclear counterstaining was performed with DAPI (Table 2). After washing (3 × 10 min in PBS), tissues were stored in PBS at 4°C in the dark until imaging using a confocal microscope (Leica SP8). Tissues were mounted serosal side up on glass slides with PBS and cover-slipped for imaging. Confocal images were acquired from the myenteric plexus plane within the muscularis externa, and only myenteric ganglia were included for neuronal quantification.

## List of abbreviations

Ach: Acetylcholine
CAIP: Cholinergic Anti-Inflammatory Pathway
EPEC: Enteropathogenic *E. coli*
VNS: Vagus Nerve Stimulation

## Acknowledgments

These studies were funded by NIH NIAID R01AI150647, R56AI179656 (C.R.). ANNA-1 antibody for neuronal staining was kindly gifted by Dr. Vanda Lennon (Mayo Clinic).

## Author contributions

K.S., E.T. and C.R. designed research; K.S., E.T., J.P., G.P., A.W., S.L., and J.L. performed research; K.S., E.T., J.P. and C.R. analyzed data; C.R. obtained funding; and K.S. and C.R. wrote the paper.

---

**Figure S1. (A)** mRNA of colon in sham, LPS, and LPS+VNS-treated mice for *Il10, Alox15, Ptsg1,* and *Ptsg2*. Data are presented as mean ± SD. One-way ANOVA and post-hoc analysis with Tukey’s multiple comparisons test were used for statistical analysis.

