## Supplementary figures and images for "Vagus nerve stimulation limits colonic inflammation through distinct neuroimmune circuitry shaped by inflammatory history"

### Supplemental Fig 1

Supplemental 1

A. Colon

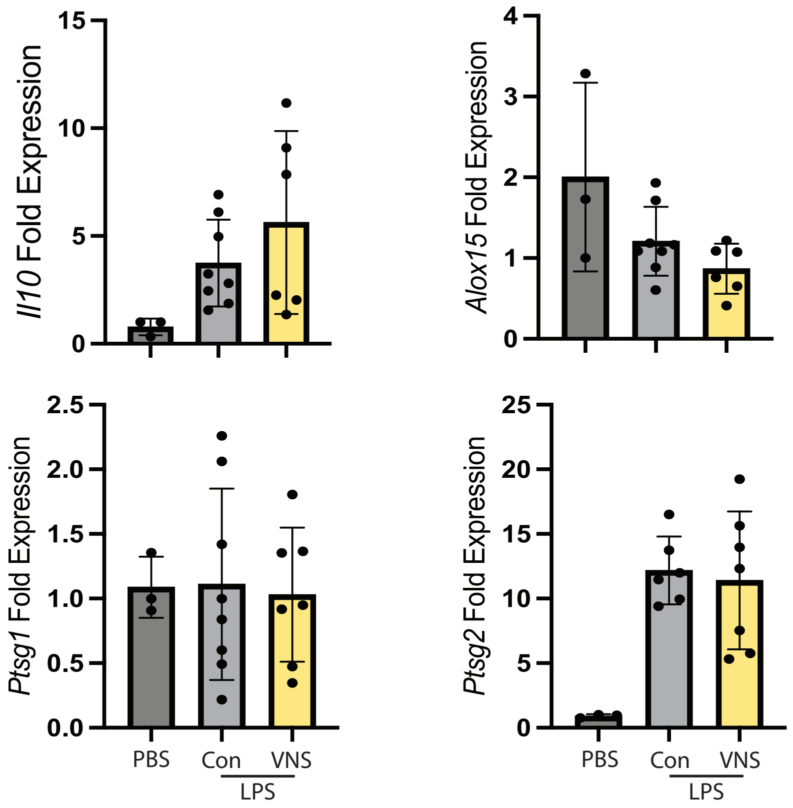
